# On Parameter Interpretability of Phenomenological-Based Semiphysical Models

**DOI:** 10.1101/446583

**Authors:** Laura Lema-Perez, Rafael Muñoz-Tamayo, Jose Garcia-Tirado, Hernan Alvarez

**Affiliations:** Universidad Nacional de Colombia, Facultad de Minas, Escuela de Procesos y Energía. Kalman research group, Cra 80 No 65-223, 050041, Medellín - Colombia; UMR Modélisation Systémique Appliquée aux Ruminants, INRA, AgroParisTech, Université Paris-Saclay, 75005, Paris, France; Center for Diabetes Technology, University of Virginia, Charlottesville, VA.

**Keywords:** Biological systems, identifiability, mechanistic models, parameter interpretability, phenomenological based semi-physical model (PBSM)

## Abstract

Empirical and phenomenological based models are used to represent biological and physiological processes. Phenomenological models are derived from the knowledge of the mechanisms that underlie the behaviour of the system under study, while empirical models are derived from analysis of data to quantify relationships between variables of interest. For studying biological systems, the phenomenological modeling approach offers the great advantage of having a structure with variables and parameters with physical meaning that enhance the interpretability of the model and its further used for decision making. The interpretability property of models, however, remains a vague concept. In this study, we tackled the interpretability property for parameters of phenomenological-based models. To our knowledge, this property has not been deeply discussed, perhaps by the implicit assumption that interpretability is inherent to the phenomenological-based models. We propose a conceptual framework to address the parameter interpretability and its implications for parameter identifiability. We use as battle horse a simple but relevant model representing the enzymatic degradation of *β*–casein by a *Lactococcus lactis* bacterium.

## 1. Introduction

How can we assess the capability of a mathematical model to provide mechanistic insight on the system under study? That is, how the mathematical structure of the model translates and captures the knowledge of the phenomena taking place in the system? To what extent can we interpret mechanistically our model? In biotechnology, biology, and biomedical fields two main approaches exist to model processes of interest, namely empirical and phenomenological based modeling. Empirical based models are derived from data, while phenomenological based models are derived from knowledge about the process. In biomedical fields, phenomenological based models are more relevant than empirical based models since, in addition to prediction, their parameters and variables provide information that can be used to perform diagnosis, discriminate clinical risk groups and guide treatment for stratifying patients by disease severity [1, 2]. In spite of this, in the fields mentioned before many models have been developed from an empirical point of view by using black box modeling approaches like machine learning and fuzzy models. Machine learning models, for example, are increasingly used in the field of medicine and healthcare but there is still an inability by humans to understand how those models work and what meaning their parameters have. Some approaches have been proposed to improving the level of explanation and interpretability of such emprirical models, that is to open the black box [3]. The deployment of the above mentioned approaches encounters its first hurdle by the difficulty of formalising the definition of central concepts such as transparency, explanation, and interpretability. In the present work, we focus on the interpretability concept but applied to phenomenological-based models. Many studies propose interpretability as a means to engender trust in empirical-based models and to reach features as close as possible to humans [1, 4, 5, 6, 7] regarding decision making. In this context, Caruana, R. et al. [1] evaluated a method for rule-based learning [8] and applied generalized additive models [2, 9, 10] to real healthcare problems to get *intelligible* and accurate models, in order to predict risk prior to hospitalizations, to have a more informed decision about hospitalization, and to reduce healthcare cost by reducing hospital admissions [1]. In the same line, Lou et al. [9, 10] call *intelligible* models to those models that can be easily interpreted by users. For decision models, the interpretability concept has been ascribed to (*i*) the ability of making decisions as close as a human being will do [11, 12], and (*ii*) the ability of being understood [11, 13]. Since, decision making is favored by the understanding of how the model works, optimal decision-based models are those that provide a trade-off between the predictive accuracy and interpretability [14].

Model interpretability is a term used in various works but without an explicit definition [11, 15]. The meaning of that term is not direct because the model as a whole is a complex piece of knowledge. Therefore, the model interpretability, scarcely will be an on-off property, i.e, a model is or is not interpretable. To grade the model interpretability will be equivalent to establish a scale of interpretability. Obviously, that scale requires a metric to generate the value of interpretability for a given model. That metric is the major problem to establishing an interpretability scale. For example, two models, one with 30 parameters and the other with only 3 parameters, but both has only one of their parameters without interpretability. If an on-off approach is maintained, both models are not interpretable. If an interpretability index (*II*) is staed as: 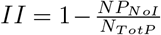 with *NP_NOI_* number of non-interpretable parameter and *N_TotP_* the total number of parameters, the *II* for first model will be 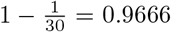 and for the second one will be 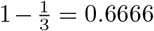. Does this proposed *II* give useful information about model size or complexity? Due to this unsolved item, in the current work the interpretability will be only evaluated in terms of individual parameters. Interpretability of model parameters is the result of multiple factors including the level of detail or specification [16], that is its granularity [17]. Due to the lacking of formalism about interpretability like a property of the parameters in a model, there is no consensus about quantifying or measuring such a property. The approach we want to elaborate in this article consists in referring the interpretability of a model to its parameters and the degree by which those parameters have physical meaning. We focus on Phenomenological Based Semi-physical Models (PBSMs) [18], of which, to the best of our knowledge, the concept of interpretability has not been deeply discussed, perhaps by the implicit assumption that interpretability is inherent to the PBSM since they are derived from a phenomenological representation of the system under study. In this work, we propose a conceptual framework that can facilitate the incorporation of interpretability for model construction. We use as battle horse a simple model to elaborate our developments. The paper is organized as follows. In Section 2, we present a summary of the steps of a modeling methodology proposed by [19] to build PBSMs. In Section 3, a conceptual framework for interpretability analysis is set using a simple mathematical model of the dynamics of enzymatic hydrolysis of *β*–casein by a *Lactococcus lactis* bacterium. Finally, we discuss in Section 4 the potential links between interpretability and identifiability. Some concluding remarks are provided in Section 5.

## 2. The process of PBSM construction

The construction of a model may be linked to a form of art. This subjective character explain the existence of several methodologies for building PBSMs [20, 21, 22, 23, 24, 25]. In our group (KALMAN, Universidad Nacional de Colombia), several studies have been developed [19, 26] to propose the following methodology, described by 10 steps, which are summarized here in the interest of completeness.

1. **Process description and model aim**: a verbal description of the process taking place is performed including a process flow diagram as graphical representation. Also, the model aim is set by the question that is expected to be answered by the model.
2. **Model hypothesis and level of detail**: a hypothesis or analogy about the behavior of the real process is proposed. Although the present methodology was originally intended for process engineering systems, it can be extended to any type of process by mean of a model hypothesis. A model hypothesis is a feasible analogy of the unknown phenomena in terms of known and well studied phenomena. If the modeled process is located in a specific area of the engineering in which the phenomena of the process are known, the hypothesis is the description of those phenomena and an analogy is not necessary. Otherwise, the process must be related to a known process, an analogy is required, and a set of assumptions is fixed. The level of detail is determined by the model objective, that is, the question that will be solved by the model.
3. **Definition of the process systems**: a process system is an abstraction of a part of the process under study [22]. Each process system (PS) is a partition of the real process, and this partition should be as real as possible, that is, physical distinctions, changes in phases or characteristics showing spatial variations in the process of interest.
4. **Application of the conservation law**: the conservation law is applied to every PS defined in step 3. Typically, mass, energy, and momentum are mainly accounted for. The equations obtained are described by either a set of ordinary differential equations in lumped models or a set of partial differential equations in distributed models; they form the basic structure of the model.
5. **Determination of the basic structure of the model**: after applying the conservation principle, select the set of equations needed to describe the model objective. Discard those equations with trivial information.
6. **Definition of the variables, structural parameters and constants**: make a list of variables, structural parameters, and constants. Variables are quantities whose values result from the solution of the model equations forming the basic structure. Parameters are values that need to be defined beforehand to solve the model. They can be known values or must be identified. Finally, the constants are fixed values either because of its universality (*e.g.*, the gravity constant) or because of the modeler choice (*e.g.*, setting a parameter with a known value from literature).
7. **Definition of constitutive and assessment equations and functional parameters**: constitutive and assessment equations are proposed to calculate the largest number of unknown parameters of each process system. The set of constitutive and assessment equations are selected according to the modeler knowledge and criteria.
8. **Verification of the degrees of freedom (DoF)**: the DoF are the difference between the number of unknowns and the number of equations.
9. **Construction of the computational model**: the solution of the mathematical model is carried out by a computational program able to solve the set of differential and algebraic equations forming the model.
10. **Model validation**: verification of the model’s domain of validity with respect to available experimental data or other validated models.

## 3. Setting a conceptual framework for interpretability analysis

In this section, we propose a conceptual framework for parameter interpretability analysis. The concepts that constitute the proposed framework to analyse parameter interpretability are defined and summarized in Table 1. For the sake of clarity, the conceptual framework is studied using a simple mathematical model that describes the dynamics of enzymatic hydrolysis of *β*-casein by a *Lactococcus lactis* bacterium in a batch system [27]. The basic structure of the model is obtained from applying a component mass balance, which results in the following unique differential equation:

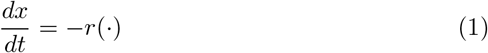

where *x* (in *μ*M) is the concentration of the substrate and *r*(⋅) (*μ*M/min) is the reaction rate, using the symbol (⋅) to indicate the dependency of this structural parameter with respect to time and any other variable or parameter of the model. It is worth to point out that global mass balance is worthless in this type of processes since no continuous inflow or outflow occurs. From Table 1, *x* is the **variable** whose dynamic trajectory is obtained by solving the model and *r*(⋅) is the unique **structural parameter**. Note that at this level of detail, the mathematical equation that represent *r*(⋅) is not yet defined. This fact suggests that for this example, Equation (1) is a unique representation of the phenomena of interest (*i.e*, the hydrolysis of *β*-casein).

**Table 1:**
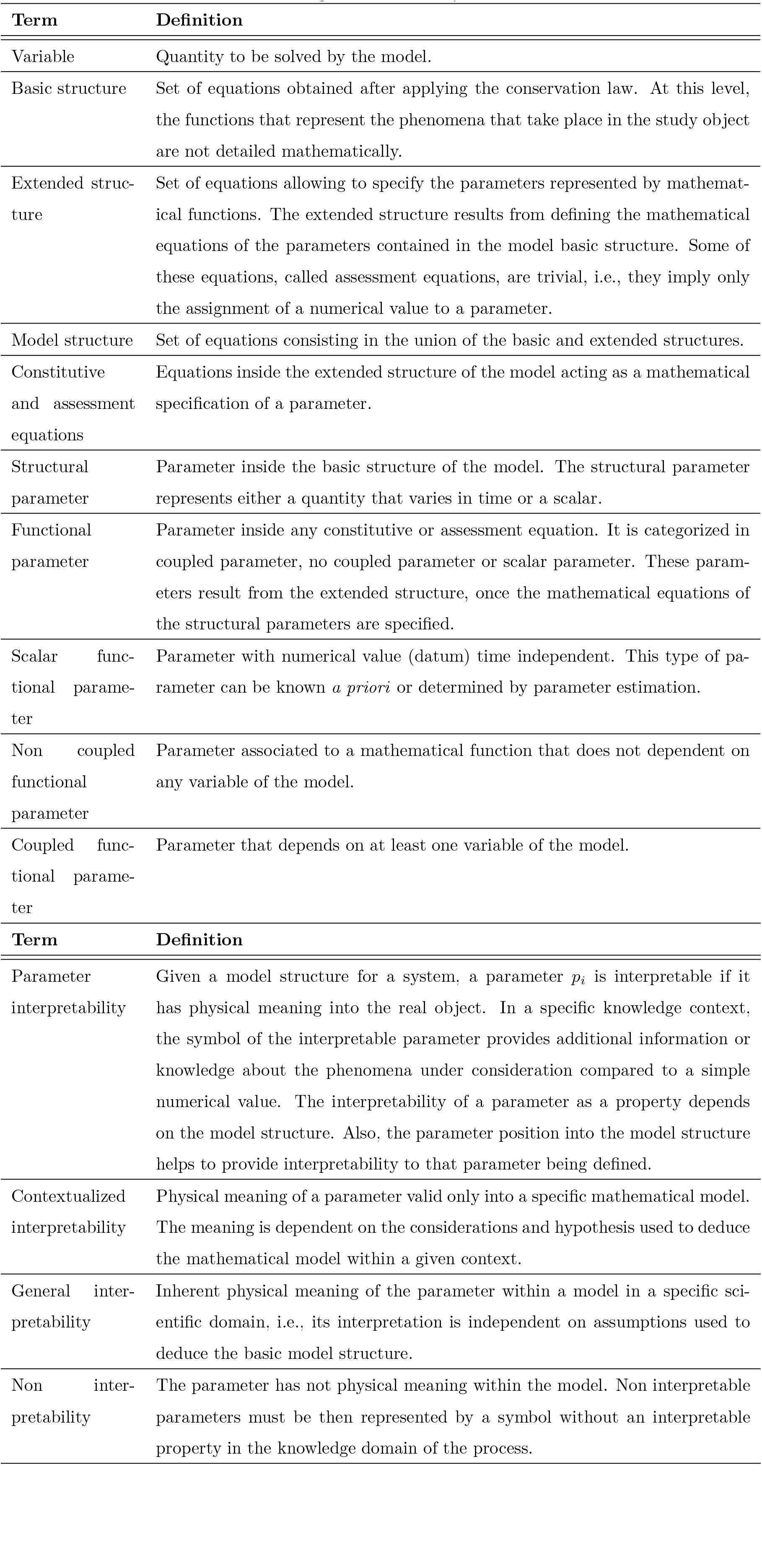
Definition of concepts used in this study

| Term | Definition |
| --- | --- |
| Variable | Quantity to be solved by the model. |
| Basic structure | Set of equations obtained after applying the conservation law. At this level, the functions that represent the phenomena that take place in the study object are not detailed mathematically. |
| Extended structure | Set of equations allowing to specify the parameters represented by mathematical functions. The extended structure results from defining the mathematical equations of the parameters contained in the model basic structure. Some of these equations, called assessment equations, are trivial, i.e., they imply only the assignment of a numerical value to a parameter. |
| Model structure | Set of equations consisting in the union of the basic and extended structures. |
| Constitutive and assessment equations | Equations inside the extended structure of the model acting as a mathematical specification of a parameter. |
| Structural parameter | Parameter inside the basic structure of the model. The structural parameter represents either a quantity that varies in time or a scalar. |
| Functional parameter | Parameter inside any constitutive or assessment equation. It is categorized in coupled parameter, no coupled parameter or scalar parameter. These parameters result from the extended structure, once the mathematical equations of the structural parameters are specified. |
| Scalar functional parameter | Parameter with numerical value (datum) time independent. This type of parameter can be known *a priori* or determined by parameter estimation. |
| Non coupled functional parameter | Parameter associated to a mathematical function that does not depend on any variable of the model. |
| Coupled functional parameter | Parameter that depends on at least one variable of the model. |
| Parameter interpretability | Given a model structure for a system, a parameter $p_i$ is interpretable if it has physical meaning into the real object. In a specific knowledge context, the symbol of the interpretable parameter provides additional information or knowledge about the phenomena under consideration compared to a simple numerical value. The interpretability of a parameter as a property depends on the model structure. Also, the parameter position into the model structure helps to provide interpretability to that parameter being defined. |
| Contextualized interpretability | Physical meaning of a parameter valid only into a specific mathematical model. The meaning is dependent on the considerations and hypothesis used to deduce the mathematical model within a given context. |
| General inter-pretability | Inherent physical meaning of the parameter within a model in a specific scientific domain, i.e., its interpretation is independent on assumptions used to deduce the basic model structure. |
| Non inter-pretability | The parameter has not physical meaning within the model. Non interpretable parameters must be then represented by a symbol without an interpretable property in the knowledge domain of the process. |

The mathematical definition of the structural parameter *r*(⋅) is the key element for the construction of the complete **model structure**, that is, for the set of equations that define the model in its basic and extended form. Multiple mathematical functions exist to define *r*(⋅) and describe the hydrolysis rate of the intact *β*-casein. In the study here analyzed [27], the authors evaluate four kinetic candidate functions to determine the best function for *r*(⋅) parameter in terms of the goodness of fit:

- First-order kinetics:

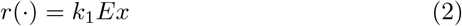
- *nth*-order kinetics:

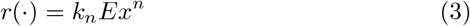
- Michaelis-Menten kinetics:

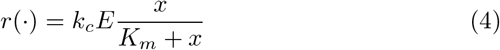
- Competitive inhibition kinetics:

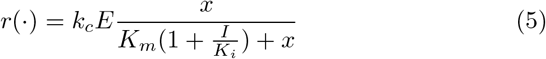

with *I* = *x*_0_ − *x*. This expression can be further manipulated to reduce the number of its parameters as:

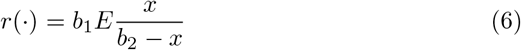

with

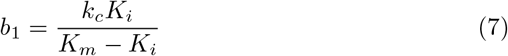

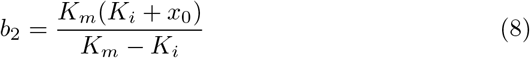

where *E* is the enzyme concentration, measured in optical density units (OD_600_). The parameter *k*_1_ (1/*OD*_600_ min) is the hydrolysis rate constant for the first-order kinetics, and *k_n_* (1/*μM*^*n*−1^*OD*_600_min) is the rate constant for the kinetics of order *n*. For the Michaelis-Menten equation, *k_c_* (*μM/OD*_600_min) denotes the catalytic rate constant and *K_m_* (*μM*) the substrate affinity constant. For the inhibition kinetics, *K_i_* (*μM*) is the inhibition constant. The concentration of the inhibitor *I* (*μM*) is considered to be equal to the concentration of *β*-casein that has been hydrolyzed (*x*_0_ − *x*), with *x*_0_ the initial protein concentration.

It is up to the modeler to decide which kinetic function to use for representing the hydrolysis rate of *β*-casein. Once, the kinetic function is defined by a new equation in addition to the basic structure, we obtain the **extended structure** of the model. The selected kinetic function is a **constitutive equation** of the model that allows to determine *r*(⋅). For example, if we select the firstorder kinetic function *r*(⋅) = *k*_1_*E_x_*, we say that *r*(⋅) is a **structural coupled parameter** that depends on the **variable** *x* and two **functional parameters**: *k*_1_ and *E*. In this case, both functional parameters have physical meaning and are thus considered to be **interpretable**. While, the enzyme concentration *E* is a known numerical value imposed by the experimental protocol, *k*1 is a rate constant that needs to be determined *via* parameter estimation.

Following the case when *r*(⋅) is specified by the first-order kinetic rate as in (2)), let’s analyze the parameter interpretability (the analysis also applies to other candidate kinetic functions, bearing in mind that the Michaelis-Menten equation is derived from a biological hypothesis on the enzyme action and thus its parameters have a stronger level of interpretability than for instance those of the kinetic of order *n*). By analyzing different experimental conditions, it was found that the hydrolysis rate of *β*-casein was dependent of the initial protein concentration *x*_0_ [27]. That is, the kinetic rate was slower at higher initial protein concentrations. To account for the dependency of the kinetic rate on the initial *β*-casein concentration, the authors performed a regression analysis with the estimated parameter values obtained for each experimental condition. After regression, the parameter *k*_1_ was further expressed as a power function of the initial *β*-casein concentration

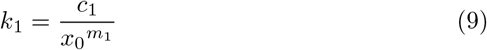

Equation (9) is referred to as a **constitutive equation**, defined by two new **functional parameters**: *c*_1_ and *m*_1_. These scalar parameters are numerical values identified by regression analysis. Table 2 shows a classification of the components of the *β*-casein model according to the conceptual framework presented in Table 1 and considering that *r*(⋅) is defined by the first-order kinetic rate in Equation (2). It is important to note that for the other kinetics options (Equations (3) - (5)) this classification is also applicable. That is, the basic structure or zero specification level is preserved, but the extended structure changes according to the chosen kinetic constitutive equation. The extended structure begins with the first specification level while the basic structure is the zero specification level and it is the only one with inherent interpretability in a PBSM.

**Table 2:**
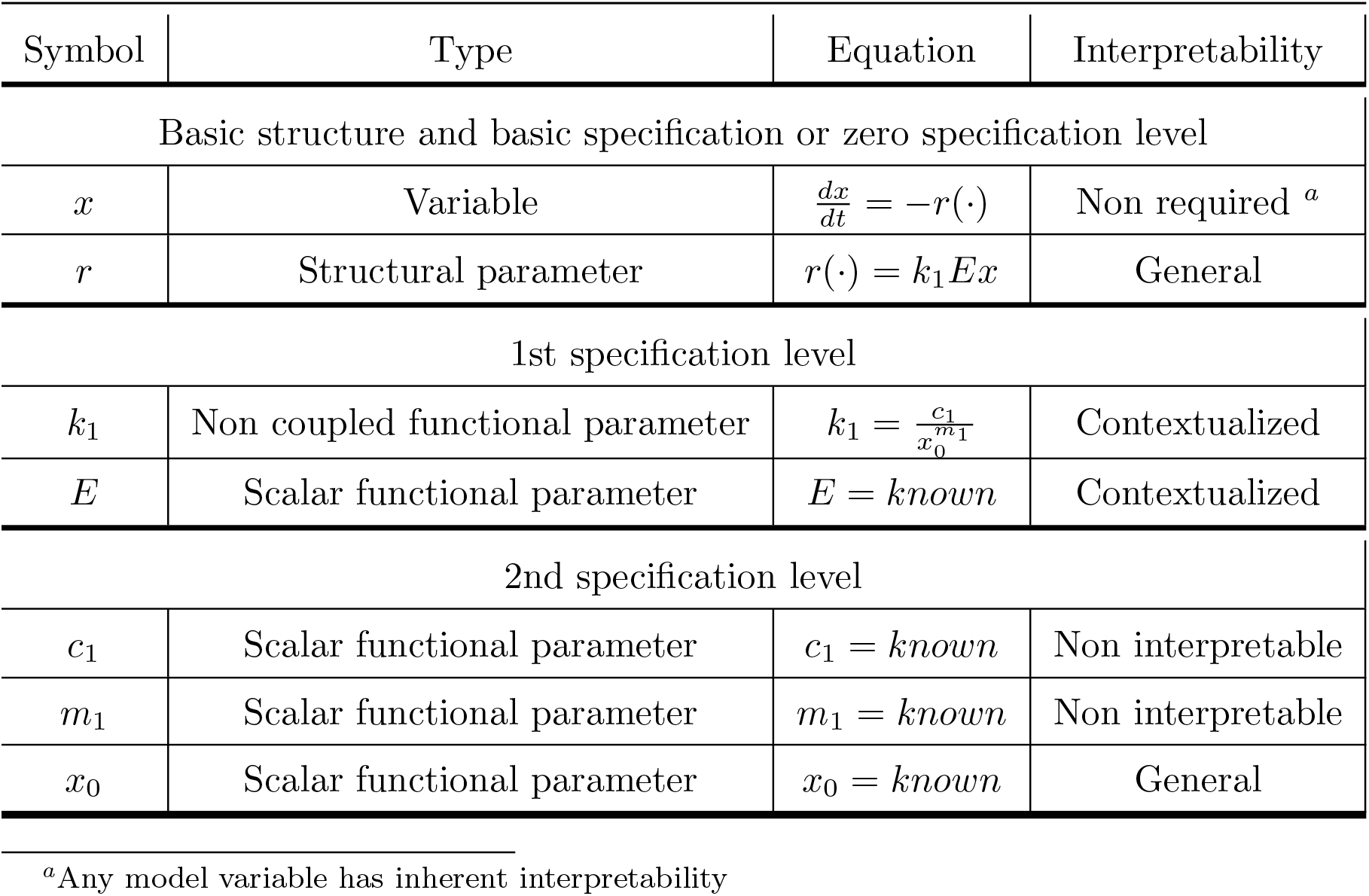
Classification of the *β*-casein model components when using the first-order kinetic rate to represent *β*-casein hydrolysis

With respect to the **parameter interpretability** of this simple model, it can be said that the **structural parameter** *r*(⋅) has **general interpretability** because in the the specific scientific domain of chemical and process engineering, the symbol *r*(⋅) denotes a reaction rate. The reaction rate determines the dynamics at which reactants are converted into products, *i.e.*, it is the number of moles of substance reacting by time unit within the reaction. The **functional parameter** *k*_1_ has **contextualized interpretability** and refers to the kinetic rate constant derived from the assumption that the hydrolysis rate follows a first-order kinetics. The **functional parameter** *E* has also **contextualized interpretability** representing the concentration of the enzyme. Contextualized means that these symbols, *k*_1_ and *E*, in other context can be used for representing another physical properties of the process.

When *k*_1_ is further defined by the constitutive equation (9) with the scalar functional parameters *c*_1_ and *m*_1_, they are **not interpretable**, since *c*_1_ and *m*_1_ are empirical parameters without physical meaning. However, the parameter *k*_1_ is still interpretable in spite of being expressed as function of non interpretable parameters. The interpretability of a parameter is not dependent on the constitutive equation that defines it in a lower specification level.

In this example, we can appreciate the peculiarity of the basic structure of a model and the dependency on the modeler choices to define the extended structure. One basic structure can lead to multiple extended structures. This extended structure results from the mathematical specification of the structural parameters. Additionally, it is highlighted how the parameters interpretability of the model can be affected when the specification levels appear, that is when the structural and functional parameters must be defined through further parametrization. A graphical explanation of the concepts applied in the example is shown in Figure 1.

**Figure 1:**
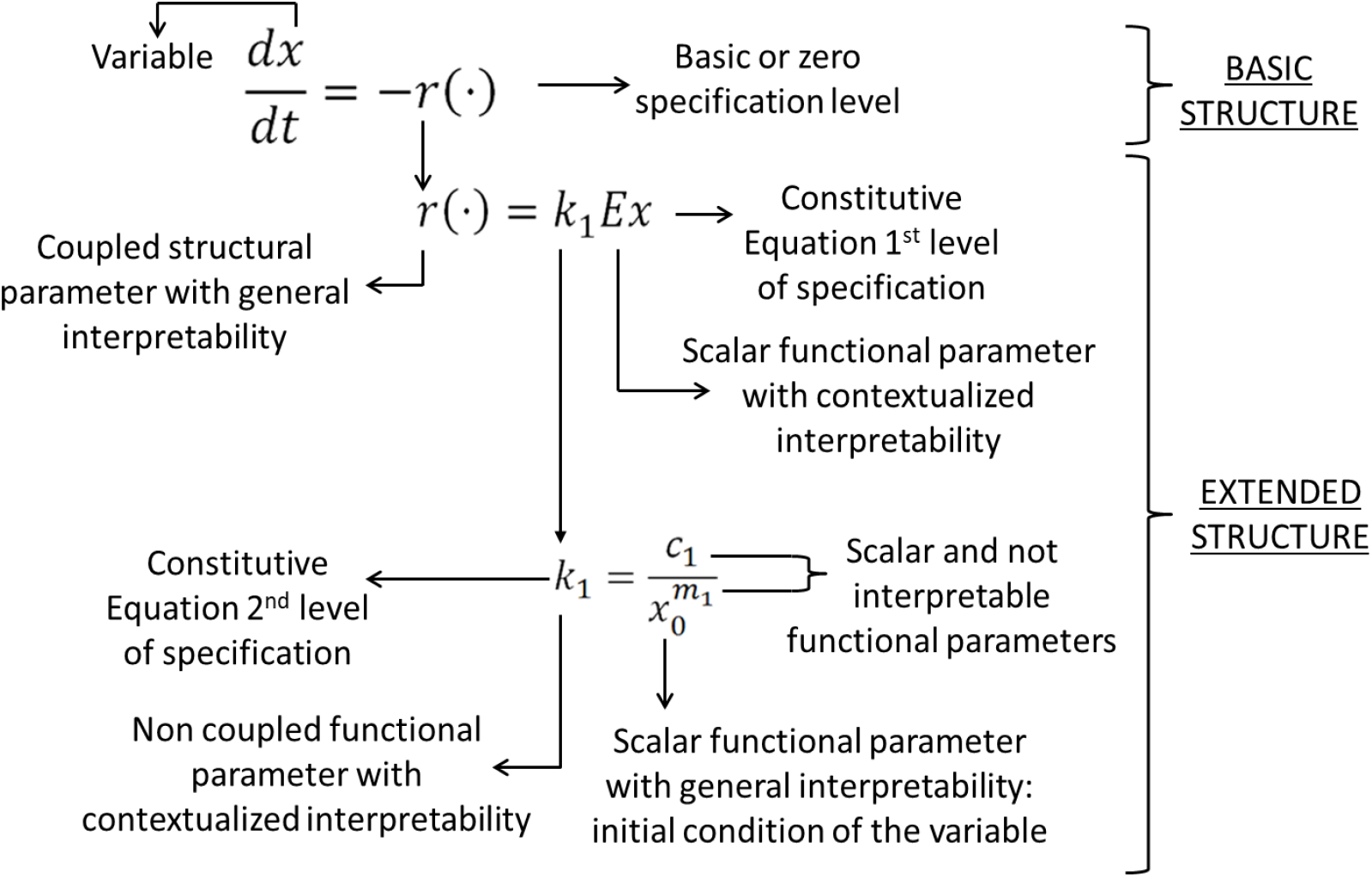
Concepts applied in a simple model of *β*-casein hydrolysis by a *Lactococcus lactis* bacterium.

## 4. Links between parameter interpretability and identifiability

In this section, we discuss about possible relations between the concepts of interpretability and identifiability.

### 4.1. Brief recall on parameter identifiability

Identifiability is a structural property of the model referred to the ability to find a unique best value of the model parameters from available measurements [28, 29]. Under the assumption that the model represents perfectly the system, model identifiability is tested in the hypothetical scenario set by continuous noise-free data and experimental conditions that provide a sufficient excitation on the model response. The structural identifiability is independent of real experimental data. Identifiability is a necessary condition for the parameter identification problem to be well posed. Identifiability testing is of great relevance for models where the parameters are biologically meaningful (as it is the case for PBSMs) and we may wish to identify them uniquely [30]. Identifiability testing can be helpful to provide guidelines to deal with non-identifiability, either providing hints on how to simplify the model structure or indicating when more information (measured data) are needed for the specific experiment [31].

Let us consider **M**(**P**) a fixed model structure with a set of parameters **P** describing the input-output behavior of the system under study. The structural identfiability of the parameter *p_i_* is determined from the following equality

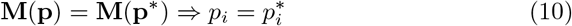

If the equality (10) holds for a unique value of the parameter *p_i_*, the parameter is structurally globally identifiable. If there are a finite number of values for *p_i_* that hold the equality (10), the parameter is structurally locally identifiable. If infinite solutions exist for *p_i_*, the parameter is nonidentifiable. A model is structurally globally (or locally) identifiable if all its parameters are structurally globally (or locally) identifiable. A model is non-identifiable if at least one of its parameters is non-identifiable. Different methods have been proposed to test identifiability of linear and nonlinear models. The interested reader is reffered to dedicated literature [32, 28, 33]. To facilitate identifiability testing, software tools such as DAISY (Differential Algebra for Identifiability of SYstems) [31] and GenSSI have been developed [34]. DAISY is implemented in the symbolic language REDUCE and GenSSI is implemented in Matlab. Both of them are freely available. We made use of both toolboxes four our analysis.

### 4.2. Interpretability vs. identifiability

In our conceptual framework, interpretability is defined as the ability to find a physical meaning of a parameter when the model structure (basic plus extended) and some knowledge of the real process are given. Interpretability is the property of the model parameters, inherited from the model structure, assigning a physical meaning to a parameter within the context where the model is constructed. When the parameter has a physical meaning, it is possible to find from available knowledge a span of numerical values to make easier its identification.

The main role of parameter interpretability for parameter identification is to narrow the search space/domain of the cost function where the identification procedure operates, constraining the values of feasible parameters to match with the existing body of knowledge. On the other hand, structural identifiability is considered a theorerical property. In practice, however, model structure misspecification and noise data can affect the identifiability of the parameters of the model [31] and therefore an accurate identification of the model parameters is not guaranteed. Practical identifiability is then subjected to the quality of available data. Interpretability can be of help in parameter identification [35] by adding prior knowledge that can be used to constraint the parameter estimation. For instance, if a parameter is interpretable, it is possible to know the threshold in which it should be placed. Also, the threshold could be restricted to improve the practical identification. A parameter can be non-identifiable, but if it is interpretable, then the prior information can be used to facilitate its practical identifiability.

Identifiability and interpretability are relevant properties of PBSMs constructed to gain mechanistic insight of the system under study. A PBSM has a basic structure that is universal and interpretable, that is, all its structural parameters are interpretable. However, it is often required to specify the structural parameters in the extended structure, yet maintaining the interpretability of a model become more challenging.

Identifiability analysis applies only to scalar parameters (see definition of scalar parameters in Table 1). In the *β*-casein model, the structural parameter *r*(⋅) is a time variant quantity and thus identifiability testing is not relevant. The quantity *r*(⋅) is interpretable and we might wonder if it is possible to estimate it from the available measurements (*x*). The reconstruction of *r*(⋅) belongs to another subject namely observability, which is not detailed here.

A structural identifiability analysis was performed for the *β*-casein model by using both DAISY software tool [31] and GenSSI-Matlab [34], to evaluate how the identifiability properties of the model change with respect to the level of specification or granularity and the candidate constitutive equations. Table 3 summarizes the identifiability and intepretability analysis. It can be noted that the basic structure of the model is interpretable but its identifiability cannot be tested because *r*(⋅) is not a scalar. However, its identifiability analysis is latter applied and is affected when the structural parameter *r*(⋅) is defined by the different kinetics. When *r*(⋅) is replaced by the first-order kinetic, the model is still identifiable. But, when *k*_1_ is further defined by a mathematical expression dependent on the initial concentration of the protein (located in the second specification level), its identifiability is modified. In the same way, for the second form of competitive inhibition kinetics, where functional parameters *b*_1_ and *b*_2_ are not replaced, the model is globally identifiable, but once *b*_1_ and *b*_2_ are defined and replaced at the next level of specification, the identifiability of the model is affected. Parameters *k*_1_, *k_n_*, *k_c_*, *K_m_*, and *K_i_* are interpretable from MichaelisMenten kinetics, but parameters *b*_1_ and *b*_2_ are not interpretable. When the mathematical expression of Michaelis-Menten is changed for the expression with parameters *b*_1_ and *b*_2_ to make easier its identification, the interpretability is affected.

**Table 3:**
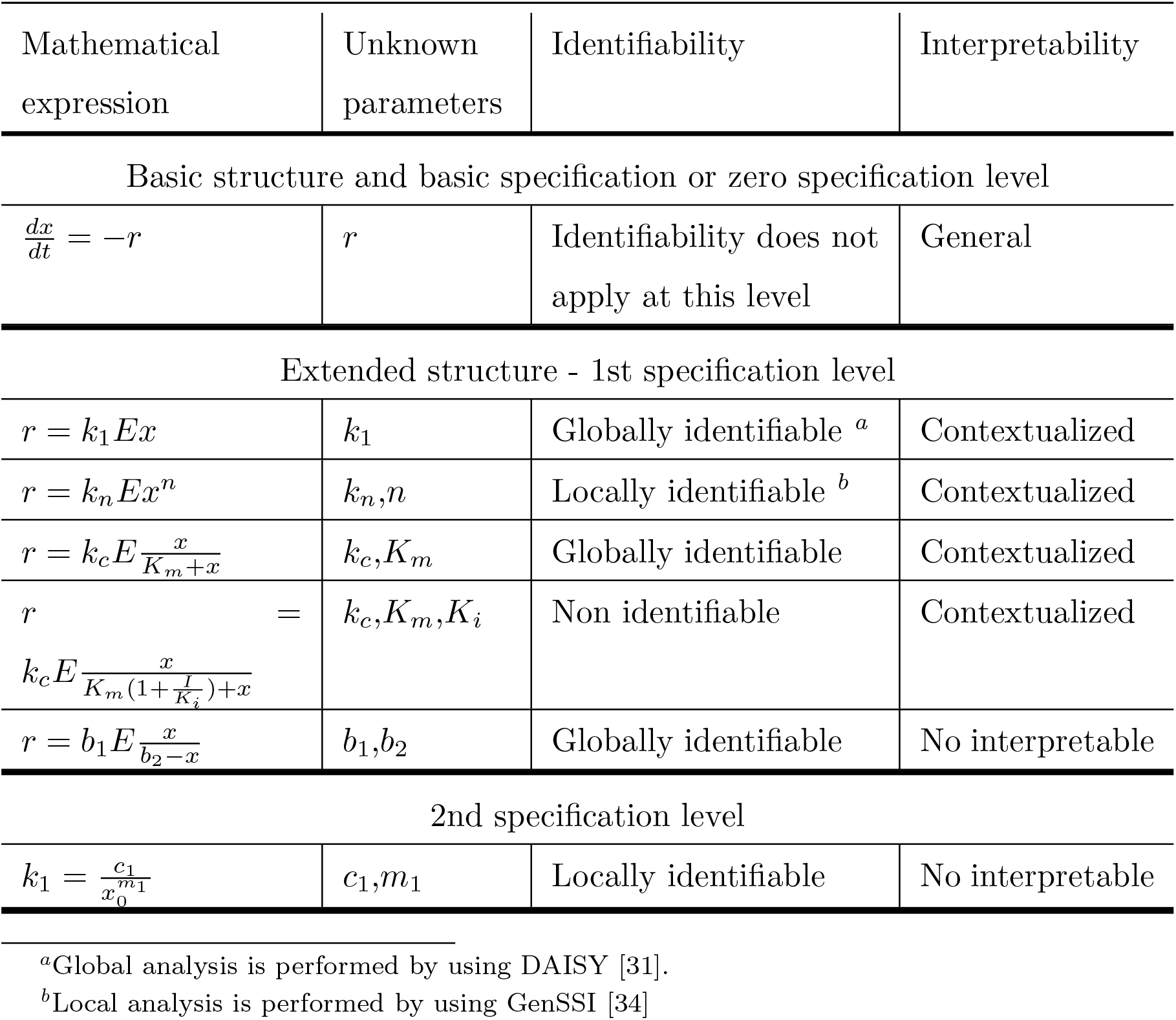
A comparison between identifiability and interpretability analysis in *β*-casein model

We deduce that a PBSM can have an extended structure to identify its parameters and an extended structure to interpret the model parameters. In the case of the *β*–casein model, two extended structures of the model can be considered depending on the interest: if the interest is to perform parameters identification, the mathematical expression containing parameters *b*_1_ and *b*_2_ is more convenient. Contrary, if the interest is to exploit the descriptive ability of the model, the mathematical expression with interpretable parameters is then selected. Note that to perform an identifiability analysis of the whole model, all parameters must be replaced by the mathematical expression defining them, whilst interpretability analysis does not require to replace the constitutive equations in the upper specification levels.

## 5. Conclusion

Due to the lack of a formal definition of the interpretability concept in the literature and that this topic is just emerging, we propose a conceptual framework for parameters interpretability. We discussed the links between parameter interpretability and identifiability.

The concepts here described provide a useful framework to undertaking the construction of models of biological/biomedical systems where the physical meaning of the model structure is a desired property. These concepts are of particularly usefulness for modeling systems that are poorly studied and thus facilitate further exploitation of *in silico* simulation. PBSMs offer great advantages for representing biological systems as they allow to enhance model capabilities in sequential way, integrate multiscale information into the same model, and guarantee direct interpretability of model basic structure. In addition, to endow with interpretability a parameter of a PBSM is an easier task when compared with the same effort over empirical models.

